# Cortical and behavioural tracking of rhythm in music: Effects of pitch predictability, enjoyment, and expertise

**DOI:** 10.1101/2023.10.15.562351

**Authors:** Anne Keitel, Claire Pelofi, Xinyi Guan, Emily Watson, Lucy Wight, Sarah Allen, Iris Mencke, Christian Keitel, Johanna Rimmele

## Abstract

The cortical tracking of stimulus features (such as the sound envelope) is a crucial neural requisite of how we process continuous music. We here tested whether cortical tracking of the beat, typically related to rhythm processing, is modulated by pitch predictability and other top-down factors. Participants listened to tonal (high pitch predictability) and atonal (low pitch predictability) music while undergoing EEG, and we analysed their cortical tracking of the acoustic envelope. Interestingly, cortical envelope tracking was stronger while listening to atonal than tonal music, likely reflecting listeners’ violated pitch expectations. Envelope tracking was also stronger with more expertise and enjoyment. Furthermore, we analysed cortical tracking of pitch surprisal (using IDyOM) and show that listeners’ expectations for tonal and atonal music match those computed by the IDyOM model, with higher surprisal (prediction errors) for atonal than tonal music. Behaviourally, we measured participants’ ability to tap along to the beat of tonal and atonal sequences in two experiments. In both experiments, finger-tapping performance was better in the tonal than the atonal condition, indicating a positive effect of pitch predictability on behavioural rhythm processing. Cortical envelope tracking predicted tapping performance for tonal music, as did pitch surprisal tracking for atonal music, indicating that conditions of high and low predictability might impose different processing regimes. We show that cortical envelope tracking, beyond reflecting musical rhythm processing, is modulated by pitch predictability, as well as musical expertise and enjoyment. Taken together, our results show various ways in which those top-down factors impact musical rhythm processing.

## Introduction

The cortical tracking of continuous auditory stimuli, such as music and speech, has been the topic of intense investigation in the past years (Keitel *et al*., 2018; Peelle *et al*., 2013; Tierney & Kraus, 2015). Cortical tracking usually refers to the neural signal matching slow amplitude fluctuations in the acoustic signal and is quantified by neural alignment to the stimulus envelope, thought to reflect processing of the rhythmic structure (Doelling & Poeppel, 2015; Gross *et al*., 2013; Luo & Poeppel, 2007). Although mostly investigated in speech, recent findings suggest that processing of naturalistic music might rely on comparable mechanisms (Harding *et al*., 2019; Sammler, 2020; Zuk *et al*., 2021). Cortical tracking is influenced by numerous top-down factors, but their interaction and relative importance are poorly understood. For example, increased attention and listening effort generally leads to stronger speech tracking (Ding & Simon, 2012; Lesenfants & Francart, 2020; Rimmele *et al*., 2015; Song & Iverson, 2018; Zion Golumbic *et al*., 2013). Conversely, both speech and music tracking are enhanced with language and music proficiency and prior knowledge (Blanco-Elorrieta *et al*., 2020; Cervantes Constantino & Simon, 2018; Di Liberto *et al*., 2018; Doelling & Poeppel, 2015; Harding *et al*., 2019). Other factors, such as the influence of enjoyment on the cortical tracking of music, have also recently elicited researchers’ interest (Weineck *et al*., 2022). Overall, any study of cortical tracking of rhythmic stimuli, needs to take into account stimuli and listener characteristics, which is one driver of the present study.

Recent studies on the cortical tracking of music have shown that the auditory cortex tracks not only the acoustic envelope but also melodic expectations, modelled as surprisal values (Abrams *et al*., 2022; Di Liberto *et al*., 2020a; Kern *et al*., 2022; Marion *et al*., 2021). These studies suggest that humans automatically process melodic expectations while listening to naturalistic, continuous stimuli (Pearce *et al*., 2010). Here, we examine the cortical tracking of pitch surprisal, using music stimuli with different levels of pitch predictability, namely tonal and atonal music excerpts. Music that is composed according to (Western) tonal principles, has an intrinsic hierarchical pitch organisation (Lerdahl, 2019). Therefore, this compositional style results in far more predictable pitch sequences than atonal music (Mencke *et al*., 2018), which is based on the compositional principle that all twelve tones within an octave are equiprobable. The few studies that have been conducted using atonal music show that the resulting lack of a hierarchical pitch organisation negatively affects memorisation (Schulze *et al*., 2011), recognition (Cuddy *et al*., 1981; Dibben, 1994; Dowling *et al*., 1995) and the strength of melodic expectations (Ockelford & Sergeant, 2013) (for review, Mencke *et al*., 2022; Mencke *et al*., 2018; Vuvan *et al*., 2014). Electrophysiological research suggests that weaker expectancies in atonal music particularly affect later attention-related processing stages (Mencke *et al*., 2021; Neuloh & Curio, 2004). Taken together, atonal music seems to present specific perceptual challenges to listeners, in particular related to melodic expectations.

In the context of musical rhythm perception, finger-tapping is often used as ‘behavioural tracking’ measure (Harding *et al*., 2019; Sammler, 2020) to assess rhythm skills (Fiveash *et al*., 2022; Iversen *et al*., 2015; Repp, 2005). The present study addresses the little-known relationship between behavioural tracking (measured by finger-tapping) and cortical envelope tracking (measured by electroencephalogram [EEG] recordings of listening participants) of musical rhythm in the context of varying pitch predictability. While temporal predictability has been shown to increase pitch discrimination performance (Herbst & Obleser, 2019; Jones *et al*., 2002), it is currently unclear whether pitch predictability affects the ability to behaviourally track naturalistic musical rhythms.

In sum, we here investigated whether cortical tracking of the music envelope, beyond rhythm processing, is modulated by pitch surprisal in two continuous, naturalistic stimulus conditions: tonal (high pitch predictability) and atonal (low pitch predictability) music. In the main experiment, we analysed participants’ EEG during passive listening, focusing on cortical envelope and surprisal tracking. We also investigated the role of enjoyment and musical expertise for cortical envelope tracking. In both the main and follow-up replication experiment, we used a behavioural measure of rhythm perception (finger-tapping) to analyse whether cortical tracking is behaviourally relevant and whether pitch predictability influences behavioural tracking. We expected that high pitch predictability in the tonal condition would be associated with better behavioural rhythm tracking than low pitch predictability in the atonal condition (see pre-registration: https://osf.io/qctpj). Due to complex and opposing effects of attention and previous experience (e.g., Reetzke *et al*., 2021) we did not hypothesise a priori on whether cortical envelope tracking would be stronger in the tonal or atonal condition.

## Materials and methods

### Participants

Twenty volunteers participated in the main study (14 female, 6 male; 18 to 26 years old; *M* = 20.95, *SD* = 1.88). It was initially planned to test 24 participants (preregistration: https://osf.io/qctpj), but data collection had to be halted due to the COVID-19 pandemic. However, the sample size analysis was based on a previous study (Schulze *et al*., 2011); *d* = .64, *α* = .05, *β* = .80; see preregistration) and yielded a desired sample size of *N* = 21, which was close to being achieved. In addition, we tested further 52 participants in a behavioural follow up experiment (see below). Participants in the main study were right-handed (*N* = 19) or ambidextrous (N = 1; Oldfield, 1971). Self-reports (Quick Hearing Check, Koike *et al*., 1994) indicated that 19 participants had normal hearing, while one reported a score that might suggest slightly diminished hearing (score of 27/60, hearing test recommended from score 20). All participants reported never having received a diagnosis of neurological/psychological disorders or dyslexia. Participants rated their musical expertise on a scale from 1 to 3 (‘none’, ‘some’, ‘a great deal’; *M* = 1.95, *SD* = 0.76). Six participants reported no musical expertise. Most participants (*N* = 18) were unfamiliar with the musical stimuli, and although two reported familiarity with the music, they could not name the piece nor composer.

The study was approved by the School of Social Sciences Research Ethics Committee at the University of Dundee (approval number: UoD-SoSS-PSY-UG-2019-84) and adhered to the guidelines for the treatment of human participants in the Declaration of Helsinki. Participants were reimbursed for their time with £15.

### Musical stimuli

Tonal and atonal polyphonic piano stimuli were used (see **Figure 1** top). For the tonal condition, we used an excerpt from W. A. Mozart’s *‘Sonata No. 5 in G Major, K. 283’*. The excerpt was taken from the second movement (*II. Andante*). The atonal piece was a manipulated version of this excerpt, created by randomly shifting the pitch of each note from 1 to 9 semitones up or down the musical scale (using GuitarPro v7.5) corresponding to 100 – 900 cents. Therefore, notes no longer formed harmonic relationships, while the timing of each note remained the same. Overall, the music in both conditions contained identical timbre, velocity, and rhythm. Each excerpt was approximately five-minutes long (292 seconds) and had a standard 4/4 time signature. The tempo of the pieces was 46 beats per minute (bpm), but because eighth note measures were consistently used, the dominant beat was 92 bpm (**Figure 1**). This equalled a rate of 1.52 Hz (see modulation spectrum in **Figure 2D**), and the beats were 652 ms apart. For the finger-tapping task, unique two-bar segments from the same pieces were extracted per condition (18 segments, each 10.4 seconds long). All music pieces were presented at a sampling rate of 44,100 Hz. All stimuli are available on the OSF server (https://osf.io/3gf6k/).

**Figure 1.**
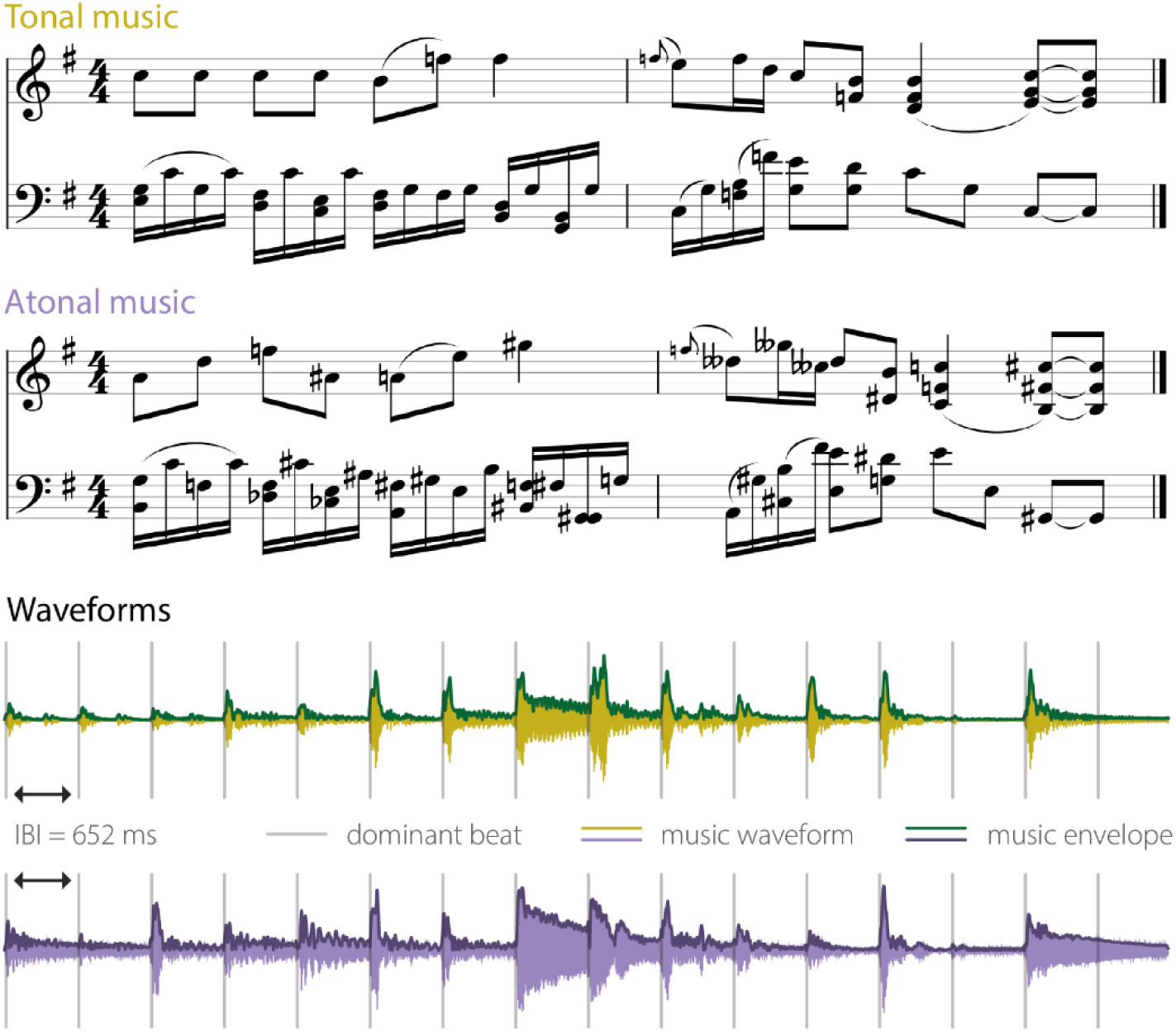
Examples of sheet music and waveforms. *Top:* Sheet music for two bars of the tonal condition (original music: *Mozart’s Sonata No. 5 in G Major, K. 283: II. Andante*) and the same bars in the atonal condition. *Bottom:* Waveforms of the same bars in the tonal (green) and atonal (purple) condition, including the music envelope. Grey bars represent the positions of the dominant beat with an inter-beat-interval of 652 ms (1.52 Hz).

**Figure 2.**
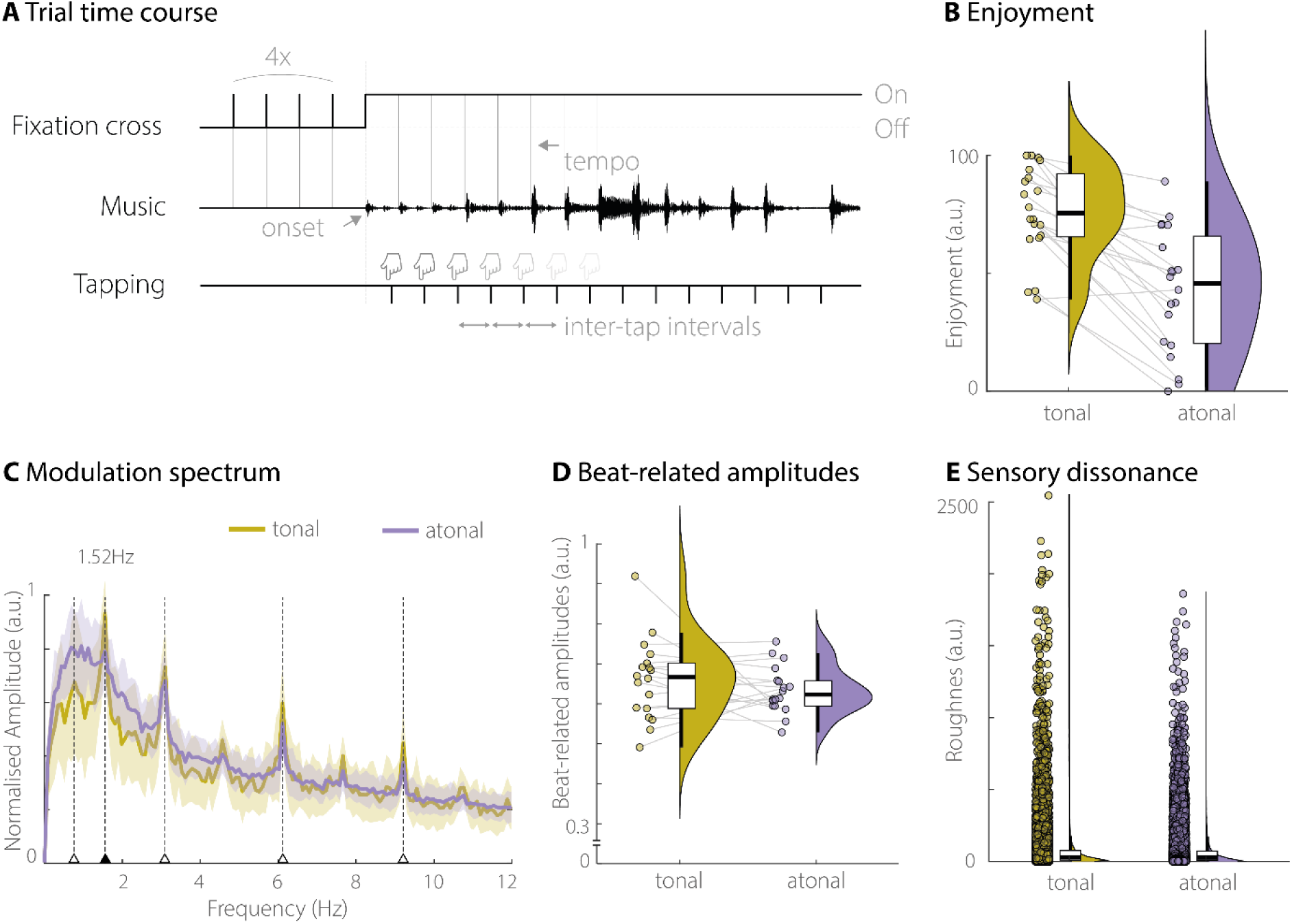
Behavioural paradigm and stimulus properties. **A)** Depiction of the trial time-course for the behavioural tracking task. Before the music started for each 2-bar trial, the dominant beat frequency (1.52 Hz) was indicated visually by flashing a fixation cross four times at that frequency. Participants tapped their finger to the dominant beat of the music once the music started. **B)** Enjoyment ratings for both 5-min tonal and atonal excerpts by all participants in the main experiment (N = 20). Overall, participants rated the tonal condition as more pleasant/enjoyable than the atonal condition. **C)** Modulation spectrum of both tonal and atonal excerpts. Thick lines indicate average values across 2-bar segments, with shaded areas representing standard error of the mean. Beat/meter related frequencies are indicated by arrows and dotted lines. **D)** Averaged amplitude values of beat-related frequencies (as shown in C) for both tonal and atonal excerpts. **E)** Sensory dissonance, assessed via roughness. There was a small difference in that the tonal condition showed larger roughness values than the atonal condition. *Notes:* Distribution plots show individual data points, box plots (including median, interquartile ranges and minimum/maximum), and kernel density estimates.

### Procedure and task

Participants performed the EEG experiment in a quiet room. They sat comfortably, approximately 110 cm from a ‘Benq’ computer screen (22.65 × 13.39 inches; 1920 × 1080 pixels resolution). On-screen instructions were presented in black, size 30 Consolas font, and displayed against a grey background. Participants could adjust the volume of the sound to a comfortable level before the start of the experimental blocks. Musical stimuli were presented using E-Prime 3.0 software (Psychology Software Tools Inc., 2016), and listened to through high-quality wired headphones (Creative, Draco HS880). Participants first passively listened to the 5-min tonal and atonal music excerpts (randomised order). Participants started the music self-paced. A five-second countdown was shown, before an ‘X’ appeared at music onset, on which participants fixated throughout the music listening. After each music piece, participants rated how pleasant they had found the music. We used a Visual Analog Scale, on which participants could rate their enjoyment by drawing a vertical line between ‘*Not pleasant*’ and ‘*Very pleasant*’.

After the passive listening blocks, participants performed a finger-tapping task to measure behavioural rhythm tracking in the tonal and atonal conditions. 36 unique trials (18 per condition) were presented in four blocks (two tonal and two atonal) of nine trials each. The order of blocks and of trials within each block was randomised across participants. Each trial was started self-paced and began with a visual presentation of the dominant beat (i.e., eighth notes). For this, an ‘X’ flashed four times at the beat frequency before the music started (see **Figure 2A**). The dominant beat was presented visually and not acoustically, to not interfere with music processing. Participants then tapped their right index finger on the outer ‘Enter’ key of a silent keyboard to the dominant beat of the music. The length of the music segments (2 bars each) required 16 finger taps per trial, resulting in approximately 288 taps per condition.

### Replication of behavioural results

To make sure that the behavioural effect found in the main experiment (more consistent finger-tapping to tonal than atonal excerpts) was robust, we carried out a follow-up replication study. All experimental procedures were ethically approved by the Ethics Council of the Max Planck Society (Nr. 2017_12). The number of participants was *N* = 52 (33 female, 19 male), and their age ranged between 20 and 41 years (*M* = 26.6, *SD* = 5.3 years). Most participants were right-handed (*N* = 43), some were left-handed (*N* = 6) or ambidextrous (*N* = 3). The procedure was identical to the main experiment, with the exception that 4 bars were used for each trial, thus doubling the time for finger-tapping per trial. This led to 10 unique tonal and 10 unique atonal trials, each 20.9 seconds long. Each trial required 32 finger taps, resulting in 320 taps per condition. Furthermore, all tonal and atonal trials were presented in random order (in contrast to tonal and atonal blocks as in the main experiment).

### Analysis of behavioural data

The inter-tap-intervals of participants’ keyboard taps for each trial were pre-processed in several ways to clean the data. First, the first two finger taps (i.e., before 981 ms) at the beginning of the trial were excluded from further analysis, to allow participants to hear two eighth notes to inform their tapping. Trials with fewer than 50% of expected remaining inter-tap-intervals were excluded (i.e., 6 necessary inter-tap-intervals in the original experiment, and 15 in the replication experiment). Inter-tap-intervals of faster than 50 ms (indicating involuntary movements) and slower than 3000 ms (indicating idling) were removed. Within each participant and condition, trials with intervals of larger than 3 standard deviations from the mean were also excluded (Abel *et al*., 2009; Rovetti *et al*., 2022; Zarate *et al*., 2015). At the participant level, our criterion was to exclude outlier data of larger than 3 standard deviations from the mean per condition (*N* = 0 in the original experiment, *N* = 0 in the replication). During the replication experiment, three participants misunderstood instructions and consistently tapped to fast 16th notes. These three participants were excluded, resulting in 49 participants that were included in the final analyses. Finger-tapping performance per trial, condition, and participant was quantified as the Median Absolute Deviation (MAD), a robust measure of dispersion (Leys *et al*., 2013) that precisely captures the variability in tapping timing. As the MAD is based on median values, it is less affected by outliers than measures based on the mean, such as variance or the coefficient of variation (Arachchige *et al*., 2022). Enjoyment ratings on the visual analogue scales to both the tonal and atonal excerpts were analysed on a scale between 0 and 100, in increments of 1 (a.u., see **Figure 2B**). We also preregistered analysing participants’ tapping accuracy in addition to their variability (https://osf.io/qctpj) but were unable to carry this out after the experiment had concluded due to restrictions on access to the laboratory and to the equipment for necessary latency measurements, as imposed by the start of the COVID-19 pandemic.

### EEG data acquisition and preprocessing

The electroencephalogram (EEG) was recorded from 32 scalp electrodes, using a BioSemi ActiveTwo system (sampling rate 512 Hz). Electrodes were placed according to the International 10-20 system. Electrodes with an offset of greater/less than ±20mV were adjusted. Ultimately, electrode offset was always below an absolute value of 30mV before the experiment began. Horizontal eye-movements were captured by two electro-oculographic (EOG) electrodes, placed at the outer canthus of each eye. To capture vertical eye-movements and blinks, a further two electrodes were positioned above and below the participants’ left eye.

Pre-processing of the EEG data was conducted using FieldTrip (Oostenveld *et al*., 2011) functions in MATLAB 2021a (MathWorks Inc.). For both 5-minute excerpts used during passive listening, we cut out epochs of 304 seconds (300 s stimulation time from music onset, plus additional 2-s leading and trailing windows). Data were initially re-referenced to *Cz* and bandpass filtered between 0.1 and 100 Hz (3^rd^ order Butterworth filter, forward and reverse). Data were then visually inspected using summary metrics (maximum value and z-value in each channel), and noisy channels were removed and interpolated using triangulation. A maximum of 4 channels was removed per participant (M = 2.47, *SD* = 0.91). Before ICA was conducted to identify blinks and artifacts, data were re-referenced to average reference (Bertrand et al. 1985). On average, *M* = 2.16 (*SD* = 0.69) components per participant were removed from the data.

### Music envelope pre-processing

To analyse the tracking of the music signal, we extracted the wideband music envelope. We first down-sampled each music excerpt to a sampling rate of 150 Hz (Keitel *et al*., 2018). Acoustic waveforms were then filtered into eight frequency bands (between 100 and 8,000 Hz, 3^rd^ order Butterworth filter, forward and reverse) that are equidistant on the cochlear frequency map (Smith *et al*., 2002). The signal in each of these eight frequency bands was *Hilbert*-transformed, and the magnitude extracted, before they were averaged for the wideband music envelope, which was used for further analyses.

### Pitch surprisal modelling

Surprisal during music listening refers to how expected a certain musical event is. Some note sequences are extremely prevalent across Western classical music, thus creating high expectations and low surprisal for an audience listening to them. To provide a computational account of music surprisal in the stimuli used, we rely on a model that learns statistical regularities of music (Pearce, 2018). Based on a variable-order Markov model, IDyOM (Information Dynamics Of Music; Pearce, 2005; Pearce & Wiggins, 2006) simulates listeners’ expectations while listening to music by collecting statistical structures of note sequences over *n*-orders on a training corpus set. Here, the training corpus was composed of a collection of Western folk songs (a subset of the Essen Folksong Collection containing 953 melodies), so as to accurately model surprisal for typical Western listeners (Guan *et al*., 2022; Kern *et al*., 2022). Specifically, the long-term component (LTM) of the model collects the sequence statistics over *n*-orders of the training set while the short-term component (STM) dynamically collects the local context over n-orders for each testing melody. For each note of the testing melodies, the model outputs a probability distribution of pitch obtained from merging distributions obtained by the STM and the LTM (for more details, see Pearce, 2005). By comparing the pitch ‘ground truth’ to the probability predicted by the model, a surprisal value is obtained. Formally, the surprisal (or information content) is the log-negative to the base 2 of the probability of the note. It essentially represents the expectedness of each note given the STM (e.g., the local context) and LTM (e.g., the long-term exposure to a musical style or culture). If surprisal is high, prediction error is also high, and vice versa (Pearce, 2018). The choice of IDyOM was motivated by numerous empirical evidence that it can accurately model a listener’s internal representation of musical regularities, both using neural and behavioural data (Di Liberto *et al*., 2020a; Gold *et al*., 2019; Kern *et al*., 2022; Pearce *et al*., 2010; Pearce & Wiggins, 2012). Since IDyOM in its current development only takes monophonic MIDI inputs, we reduced the complete score of each excerpt into a monophonic version that contained the melody and the bass line. The pitch surprisal values for each note were then used to build a continuous signal, with surprisal values making up the amplitude for the duration of the respective note. This initial step function was smoothed by convoluting it with a Gaussian filter (*sigma* = 50). The continuous surprisal signal was created to have the same sampling rate as the EEG signal (150 Hz).

### Mutual information analysis

The correspondence between the continuous EEG signal and envelope and surprisal signals (i.e., cortical envelope tracking and cortical surprisal tracking) was analysed using a Gaussian Copula mutual information (MI) framework (Ince *et al*., 2017; Keitel *et al*., 2017). In this approach, which is optimised for neurophysiological data, Gaussian copulas are used to normalise the continuous, analytical signals (Ince *et al*., 2017). The first 500 ms of the signals were removed from analysis, to avoid contamination with strong transient evoked responses at the start of the music. Mutual information (in bits) between the EEG signal and the music envelope was computed with both signals filtered at the dominant beat frequency range (0.5 – 2 Hz). A participant-specific stimulus-brain lag was included for the envelope tracking analyses, which was based on the individual phase coherence (Harding *et al*., 2019) peak at auditory electrode Cz, which was averaged for slow frequencies between 1 and 12 Hz (before the narrow band-pass filtering described above). Initial coherence values were computed for lags between 40 and 200 ms in steps of 20 ms.

Mutual information between the EEG signal and the pitch surprisal signals was computed with signals bandpass filtered between 0.1 and 50 Hz. As the analysis sampling rate was 150 Hz, filtering up to 50 Hz allowed for this frequency to be robustly reflected in the data (1/3 of the sampling frequency). This wide range was chosen as no clear assumptions about a specific, narrow-band frequency range could be made and prediction processes have been shown across multiple frequency bands (Engel *et al*., 2001; Morillon *et al*., 2019). The surprisal-tracking analysis included surprisal values for both the melody and bass lines jointly, capitalising on the multivariate capabilities of the used MI approach (Ince *et al*., 2017). Apart from the wider frequency band, the analysis of surprisal tracking was equivalent to that of envelope tracking, including the same individual stimulus-brain lags.

Each mutual information value was computed per participant, condition, and electrode. The results of these analyses will be referred to as cortical (envelope or surprisal) ‘tracking’, and we do not make assumptions about the underlying mechanisms (e.g., cortical entrainment) as these are still debated (Alexandrou *et al*., 2018; Keitel *et al*., 2021; Obleser & Kayser, 2019).

### Statistical analyses

To test the statistical significance of mutual information values for envelope and surprisal tracking against chance, 3000 permutations were computed per participant, condition, and electrode. Specifically, to create permuted data, we segmented the continuous envelope/surprisal signals into 1-s segments and shuffled the segments randomly. This kept the statistical properties of the signal but destroyed the temporal relationship between the music and brain signals. MI was then computed between the brain signal and the 3000 shuffled envelope/surprise signals. The group level mean was then tested against the 95th percentile of the random group mean distribution, essentially implementing a one-sided randomisation test at *p* < .05 (Brohl *et al*., 2022).

For the comparison between the two conditions, *t*-values were computed using the real MI values, as well as the 3000 MI values from the shuffled data. These real and permuted data were then compared, again using a cluster-based permutation test, with a critical *t-*value of 2.1, which represents the critical value of the *Student’s t* distribution for 20 participants and a two-tailed probability of *p* = .05 (Keitel *et al*., 2018).

Pearson’s correlations between cortical tracking and behavioural measures (tapping variability, musical competency, and enjoyment) were computed between the behavioural measures and the true MI values, as well as between the behavioural measures and the 3000 permuted MI values. Before comparing the true *r*-values with the permutation distribution using cluster-based permutation (as above), Pearson’s *r*-values were transformed to be normally distributed using Fisher’s z-transformation (e.g., Gorsuch & Lehmann, 2010). For all cluster-based permutation analyses, initial clusters were chosen at an alpha level of *p* < .05. As an indicator of effect sizes, we either report *Cohen’s d* for peak electrodes in the case of *t* or *r* values (Brohl & Kayser, 2021; Lakens, 2013), or summed MI values within each significant cluster (*MI*_sum_) (Keitel *et al*., 2018).

To be able to draw evidence-led conclusions about the laterality of our results, we explicitly tested for hemispheric lateralisation (Keitel *et al*., 2020; Park & Kayser, 2019). Hemispheric differences in cortical tracking have been theoretically assumed and occasionally been found experimentally for the music envelope (Doelling & Poeppel, 2015; Zatorre, 2001). The participant-specific results (e.g., MI values) were extracted for significant electrodes in one hemisphere and in corresponding contralateral electrodes. We then averaged these values within each hemisphere and calculated the between-hemispheres differences with a group-level, *Student’s t*-test (two-sided). *P*-values were corrected for multiple comparisons using FDR correction at the level of 5% (Benjamini & Hochberg, 1995).

All tests are *two-tailed*, except for the comparison of finger-tapping variability between the tonal and atonal condition. These comparisons are one-tailed, as we had an *a priori* directed hypothesis that finger-tapping in the atonal condition would be more variable than in the tonal condition (see preregistration, https://osf.io/qctpj).

## Results

### Differences between tonal and atonal music stimuli

As intended, the pitch surprisal was overall higher for the atonal than the tonal stimuli (see **Figure 5B**). Pitch surprisal was computed for all notes, separately for melody and bass lines using IDyOM (Pearce, 2005). The surprisal is estimated by comparing the pitch ground truth of a note to its predicted value in the model’s output distribution. For the melody line, pitch surprisal was on average *M* = 3.02 (*SD* = 3.28) in the tonal condition, and *M* = 6.96 (*SD* = 3.78) in the atonal condition (see **Figure 5B**). A *Student’s t*-test confirmed that surprisal values were statistically larger in the atonal than the tonal condition, *t*(597) = −25.21, *p* < .001, *Cohen’s d* = −1.03. Likewise, pitch surprisal in the bass line was higher in the atonal than in the tonal condition (tonal: *M* = 3.26, *SD* = 2.72; atonal: *M* = 7.38, *SD* = 3.54; *t*(722) = −28.41, *p* < .001, *Cohen’s d* = −1.06).

Furthermore, participants were asked to rate how pleasant they found listening to the tonal and atonal music stimuli (on a scale effectively analysed from 0 to 100 a.u., **Figure 2B**). The enjoyment ratings indicated that participants found the tonal excerpt more pleasant than the atonal excerpt (tonal: *M* = 76.18, *SD* = 19.31; atonal: *M* = 42.53, *SD* = 25.99; *t*(19) = 6.47, *p* < .001, *Cohen’s d* = 1.45).

We also computed the modulation spectrum (Ding *et al*., 2017) for both conditions between 0 and 12 Hz (**Figure 2C).** This showed several peaks at beat-related frequencies (i.e., subharmonic and harmonics of 1.52 Hz). A comparison of average beat-related frequencies (c.f., Celma-Miralles & Toro, 2019; Nozaradan *et al*., 2012) for excerpts across both conditions showed no statistical difference (tonal: *M* = 0.66, *SD* = 0.10; atonal: *M* = 0.63, *SD* = 0.06; *t*(17) = 1.02, *p* = .322, *Cohen’s d* = 0.24; **Figure 2D**). This indicates that the amplitudes of beat-related peaks in the modulation spectrum were comparable in both conditions.

To assess whether the atonal condition shows any drastic perceptive pitch differences compared with the tonal condition, we computed roughness as measure of sensory dissonance (using the MIR toolbox; Lartillot *et al*., 2008). The default frame length of 50 ms (25 ms overlap) was used. The roughness in the tonal condition was slightly higher than in the atonal condition (tonal: *M* = 82.85, *SD* = 155.73; atonal: *M* = 60.18, *SD* = 122.02; *t*(11674) = 15.25, *p* < .001, *Cohen’s d* = 0.14; **Figure 2E**). The effect was very small (according to *Cohen’s d* conventions) but implies at least that the atonal condition did not show more sensory dissonance than the tonal condition.

### Behavioural tracking: Finger-tapping is more variable in the atonal than the tonal condition

To test how pitch predictability influences behavioural rhythm tracking, we first analysed differences in inter-tap-intervals between the tonal and atonal conditions. To mitigate the effect of potential outliers on the group level (see **Figure 3**), we used a non-parametric approach. In the Experiment 1, the median absolute deviation (MAD) of inter-tap-intervals in the tonal condition was on average *M* = 29.72 ms (*SD* = 14.99 ms). In the atonal condition, the MAD was slightly higher, on average *M* = 40.30 ms (*SD* = 35.66 ms). A direct comparison using a Wilcoxon Signed Rank test indicated that tapping performance was significantly more variable in the atonal than the tonal condition (difference = 10.6 ms, *Z* = −1.94, *p* = 0.026, *one-tailed*). Out of the 20 participants, 15 (75%) had more variable inter-tap-intervals when tapping in the atonal than in the tonal condition.

**Figure 3.**
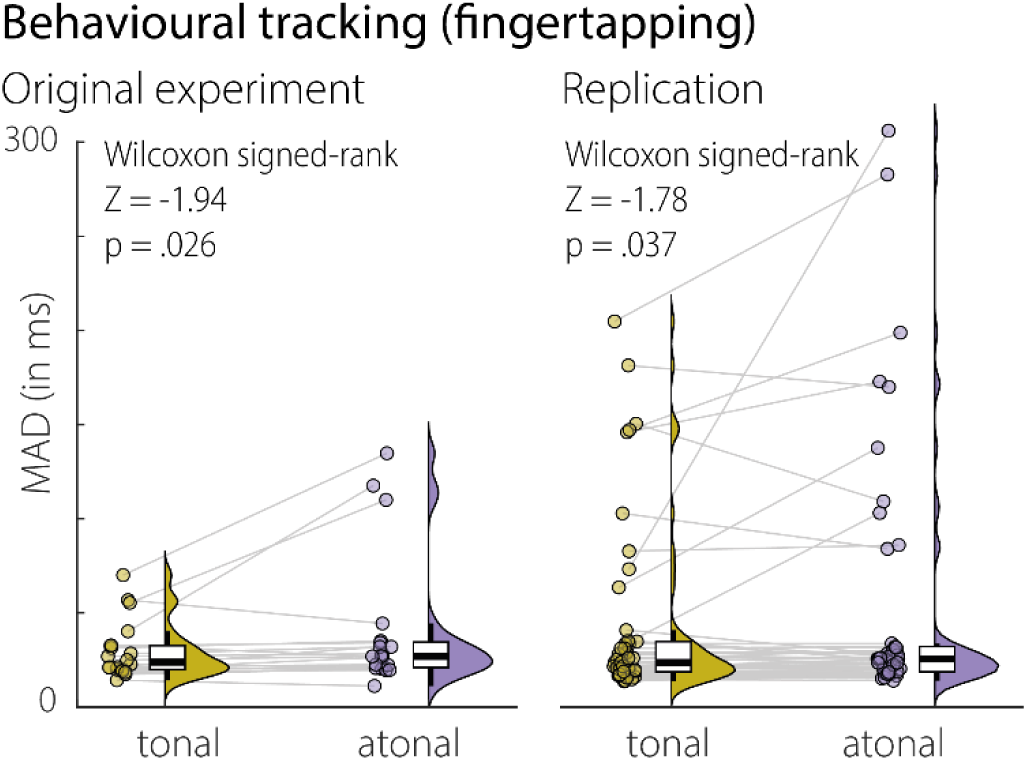
Behavioural tracking results of the main experiment (*N* = 20) and the replication experiment (*N* = 49). Shown is the Median Absolute Deviation (MAD) in ms for inter-tap-intervals in both the tonal and atonal condition. Points indicate individual data for all participants, violin plots show kernel density estimates, and boxplots show median interquartile ranges and minimum/maximum.

We performed the same analysis for the behavioural follow-up study, which had more than twice as many participants, and in which individual trials were twice as long as in the original experiment. The MAD of inter-tap-intervals in the tonal condition was on average *M* = 42.56 ms (*SD* = 46.24 ms). In the atonal condition, the MAD was again slightly higher, on average *M* = 52.42 ms (*SD* = 67.33 ms). A Wilcoxon Signed Rank test confirmed that tapping performance was significantly more variable in the atonal than the tonal condition (difference = 9.9 ms, *Z* = −1.78, *p* = 0.037, *one-tailed*). Out of the 49 participants, 30 (61.2%) had more variable inter-tap-intervals when tapping in the atonal than in the tonal condition. Together, the results of the original and replication experiments indicate that there is a small but replicable effect of tonality on finger-tapping variability: When listening to tonal music, participants tap to the beat more consistently than when listening to atonal music.

### Cortical tracking of the music envelope

We first analysed whether participants tracked the acoustic music envelope, band-pass filtered around the dominant beat frequency, in the tonal and atonal condition compared with chance level, using cluster-based permutation (**Figure 2A**). In the tonal condition, we found a large positive cluster of 31 electrodes that tracked amplitude fluctuations significantly (*p* < .001, *MI*_sum_ = 0.332). Equivalently, in the atonal condition, there was a positive cluster of 32 electrodes that showed significant envelope tracking (*p* < .001, *MI*_sum_ = 0.438). There was no evidence for a hemispheric lateralisation, neither in the tonal nor the atonal condition (both *p*_FDR_ > 0.82). We then directly compared envelope tracking in both conditions. This resulted in two negative clusters, one over frontal electrodes (*p* < .001, *Cohen’s d*_peak_ = −1.17, 7 electrodes) and one over left occipital electrodes (*p* = .003, *Cohen’s d*_peak_ = −2.09, 3 electrodes). These negative clusters indicated that the acoustic music envelope was tracked stronger in the atonal than the tonal condition.

### Envelope tracking for tonal music during passive listening predicts finger-tapping performance

To test whether envelope tracking during passive listening predicted participants’ behavioural tracking of the beat (i.e., finger-tapping performance), we correlated the MI values per electrode with participants’ average tapping variation (MAD, **Figure 4A**). We found one negative cluster over left-frontal electrodes that predicted tapping variance in the tonal condition (*p* = .026, *Cohen’s d*_peak_ = −1.02, 2 electrodes). These results indicate that participants who showed stronger envelope tracking to tonal music had smaller variance (i.e., better performance) when tapping, than participants with weaker envelope tracking. Envelope tracking when listening to atonal music did not significantly predict tapping performance in the atonal condition. To compare this relationship directly between the tonal and atonal condition, we entered the average MI values of electrodes in the negative cluster as predictor into a regression model, with *tonality* as additional predictor, an *envelope tracking × tonality* interaction term, and *finger-tapping variability* (MAD) as outcome variable. This overall model was not significant (*F*(3,26) = 1.02, *p* = .395) and only explained 7.84% of the variance. The model yielded no main effects (both *p* > .32) and no interaction (*p* > .21), which suggests that the effect of envelope tracking on finger-tapping is small in the tonal condition, and not statistically different between the tonal and atonal condition.

**Figure 4.**
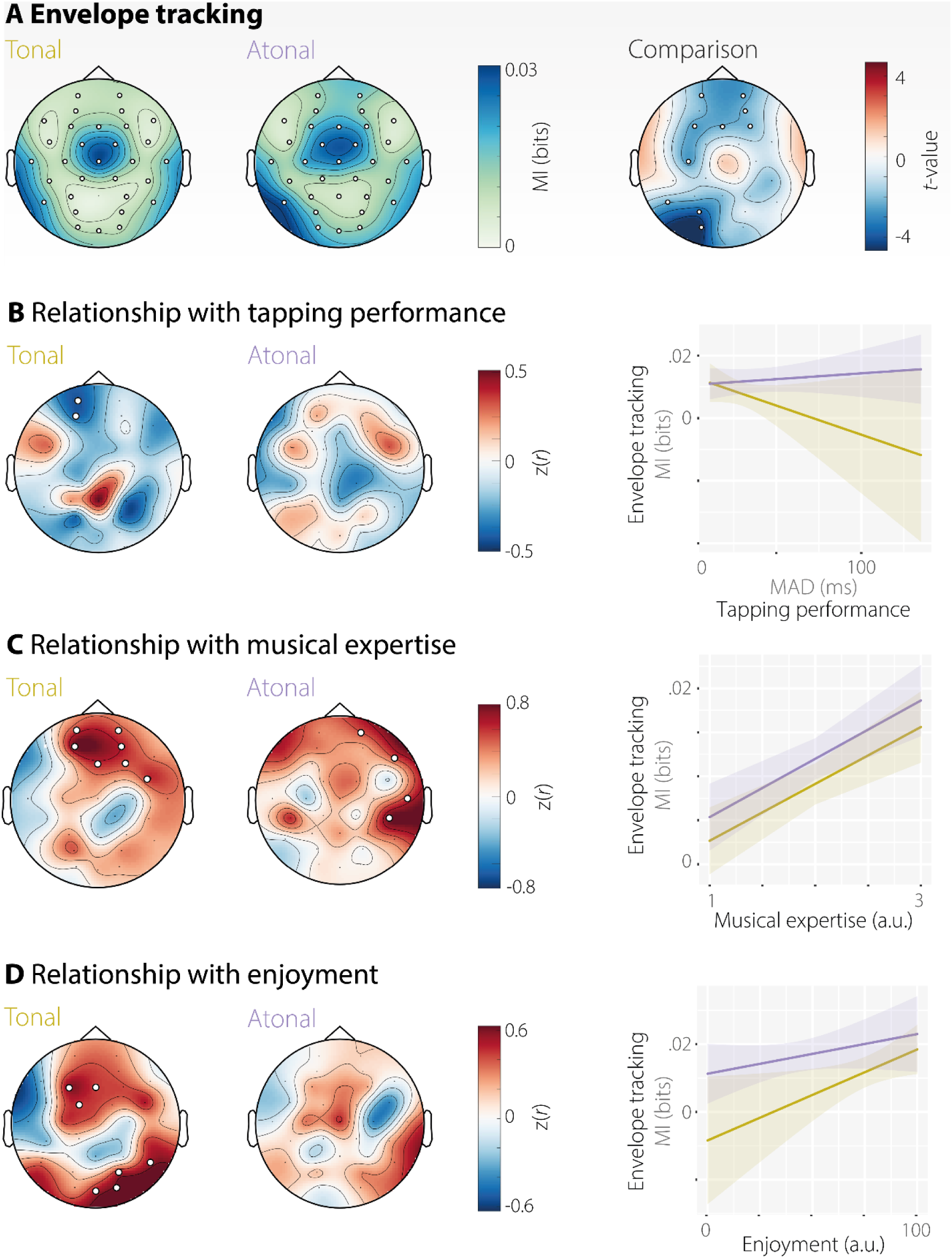
Cortical tracking of acoustic music envelope and its relationship with behavioural measures. **A)** Topography of cortical envelope tracking assessed through mutual information (in bits) for both conditions. The right topography shows *t*-values from a direct comparison between tonal and atonal music. **B)** Correlation between cortical envelope tracking and participants’ finger-tapping performance. Envelope tracking predicted finger-tapping performance only in the tonal condition, in a left-frontal cluster. Here, stronger envelope tracking was associated with better performance (i.e., less tapping variability). **C)** Correlation between cortical envelope tracking and participants’ self-reported musical expertise. Expertise predicted envelope tracking in both conditions, in a frontal (tonal) and fronto-right-lateral (atonal) cluster. Stronger envelope tracking was associated with more musical expertise. **D)** Correlation between cortical envelope tracking and enjoyment. Enjoyment predicted envelope tracking in the tonal condition in fronto-central and posterior electrodes. A regression model showed a significant main effect of enjoyment with no significant interaction (*p* = .06). Perceiving the music as more pleasant was associated with stronger envelope tracking. Note: Significant electrodes are highlighted with white circles.

### Expertise predicts envelope tracking for tonal and atonal music

Several previous studies have found that musical expertise is associated with stronger neural synchronisation to music (Di Liberto *et al*., 2020b; Doelling & Poeppel, 2015; Harding *et al*., 2019). We therefore tested the relationship between participants’ musical expertise and acoustic envelope tracking. A large fronto-temporal cluster showed a significant positive correlation between self-assessed musical competency and music tracking in the tonal condition (*p* < .001, *Cohen’s d_peak_* = 2.76, 7 electrodes). In the atonal condition, envelope tracking was also positively predicted by a fronto-temporal cluster (*p* = .004, *Cohen’s d_peak_* = 2.85, 4 electrodes). A regression model predicting *envelope tracking* (averaged across the electrodes included in the significant clusters reported above) from *tonality, musical expertise*, and their interaction, (*F*(3,26) = 12.83, *p* < .001) explained 51.7% of the variance. Only the main effect of *expertise* was significant (*t* = 4.15, *p* < .001; main effect of *tonality* and the interaction both *p* > .59). These results indicate that musical expertise is associated with enhanced tracking of the music envelope for both high predictable (tonal) and low predictable (atonal) music but is unlikely to explain differences in tracking between conditions.

### Enjoyment predicts envelope tracking for tonal music

Participants also indicated how pleasant they found listening to the music after each condition using visual analogue scales. These enjoyment ratings were significantly higher for tonal than atonal music (*Cohen’s d* = 1.45; see above and **Figure 2C**). Ratings were correlated with cortical envelope tracking across participants. In the tonal condition, two significant clusters showed a positive correlation between enjoyment and envelope tracking, one frontocentral cluster (*p* = .026, *Cohen’s d_peak_* = 1.40, 3 electrodes [F3, FC1, Fz]) and one occipital cluster (*p* = .004, *Cohen’s d* = 1.61; 4 electrodes [Oz, O2, PO4, P8]). No significant clusters emerged in the atonal condition. A regression model predicting *envelope tracking* (averaged across the electrodes included in the significant clusters reported above) from *enjoyment* ratings, *tonality* and their interaction (*F*(3,26) = 2.92, *p* = .047) explained 19.6% of the variance. More *enjoyment* increased cortical tracking (*t* = 2.28, *p* = .029), whereas neither *tonality* (*t* = 1.92, *p* = .062), nor an interaction of *tonality* with *enjoyment* affected tracking (*t* = −1.03, p = .310).

### Cortical tracking of pitch surprisal

Pitch surprisal was analysed using the IDyOM model (Pearce, 2005) in both conditions. As expected, surprisal was higher for notes in the atonal than the tonal condition for both the melody and bass lines (*Cohen’s d_peak_* = −1.03 and *Cohen’s d_peak_* = −1.06, respectively; see above and **Figure 5B**). We first analysed whether pitch surprisal was tracked in both conditions above chance level, in a multivariate analysis including surprisal in melody and bass lines. In the tonal condition, pitch surprisal was tracked in a large bilateral cluster (**Figure 5A**; *p* < .001, *MI*_sum_ = 0.073; 16 electrodes). Likewise, in the atonal condition, pitch surprisal was tracked in a bilateral electrode cluster (*p* < .001, *MI*_sum_ = 0.051; 14 electrodes). Although pitch tracking in the tonal condition appeared to be larger than in the atonal condition, directly comparing the tracking of pitch surprisal between both conditions yielded no statistically significant clusters. Likewise, although pitch surprisal tracking appeared larger in the right hemisphere, contrasting left- and right-hemispheric cluster electrodes yielded no systematic lateralisation of cortical tracking (tonal: *p*_FDR_ =.27 and atonal: *p*_FDR_ =.13, respectively). These results suggest that listeners form pitch expectations (i.e., prediction errors) comparable with the IDYOM model, and that pitch surprisal is represented in the brain to a similar extent in tonal and atonal conditions.

**Figure 5.**
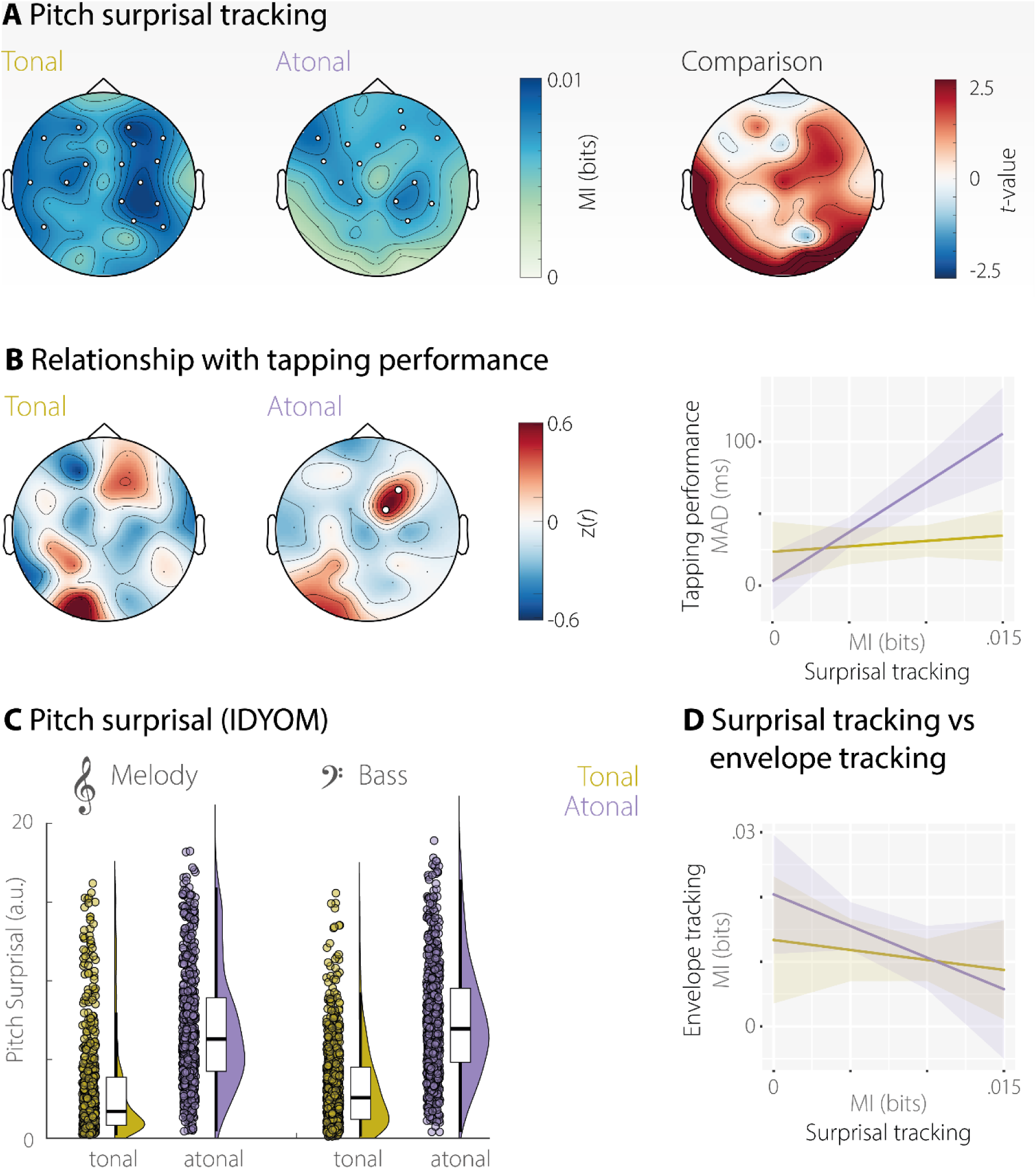
Cortical tracking of pitch surprisal performance. **A)** Topography of cortical surprisal tracking assessed through mutual information (in bits) for both conditions. The right topography shows t-values from a direct comparison between tonal and atonal music. Significant electrodes are highlighted with white circles. **B)** Correlation between pitch surprisal tracking and participants’ finger-tapping performance. Surprisal tracking predicted finger-tapping performance only in the atonal condition, in a right-frontal cluster. Here, stronger surprisal tracking was associated with worse performance (i.e., higher tapping variability). C) Pitch surprisal values for each note in the melody and bass lines of both tonal and atonal 5-min excerpts. Points indicate data for all notes, violin plots show kernel density estimates, and boxplots show median interquartile ranges and minimum/maximum. Pitch surprisal was higher for the atonal than the tonal condition. **C)** Relationship between cortical tracking of acoustic envelope and pitch surprisal.

### Surprisal tracking predicts finger-tapping performance in the atonal condition

We also analysed whether the extent to which participants tracked pitch surprisal predicted their finger-tapping performance. This correlation analysis yielded no significant clusters in the tonal condition. However, in the atonal condition, the tracking of pitch surprisal was positively correlated with finger-tapping performance in one frontocentral cluster (**Figure 5B**; *p* = .027, *Cohen’s d_peak_* = 1.11, 2 electrodes). Again, to be able to draw conclusions about differences between the tonal and atonal condition, we entered the average MI values of the positive cluster as predictor into a regression model, with *tonality* as additional predictor, a *surprisal tracking × tonality* interaction term, and *finger-tapping variability* (MAD) as outcome variable. The overall regression model (*F*(3,36) = 7.26, *p* < .001) explained 37.6% of variance in finger-tapping variability. Neither the main effect of *surprisal tracking* (*p* > .49) nor the main effect of *condition* (*p* > .16) reached significance. However, the interaction *surprisal tracking × condition* was statistically significant (*t* = 3.20, *p* < .003). This interaction stemmed from an effect, exclusive to the atonal condition, where participants who tracked the pitch surprisal well, finger-tapped with higher variability than participants with relatively poor cortical tracking.

### Relationship between envelope tracking and surprisal tracking

Last, we were interested in the relationship between acoustic envelope tracking and pitch surprisal tracking, because these measures have not previously been looked at together. We used the average MI value per cluster (as seen in **Figure 4A** and **Figure 5A&B**) and participant in a regression model with *acoustic envelope tracking* as outcome variable and *surprisal tracking*, *tonality* and a *surprisal tracking × tonality* interaction term as predictors. The overall regression model was not significant (*F*(3,36) = 1.67, *p* = .190) and explained 12.2% of the variance. No main or interaction effect reached significance (all *p*-values > .295; **Figure 5D**). Thus, there seems to be no systematic relationship between acoustic envelope tracking and pitch surprisal tracking.

## Discussion

In the present study we show that the cortical representation of naturalistic continuous music, as measured through envelope tracking, reflects not only rhythm processing, but is also modulated by pitch predictability, musical expertise, and enjoyment (**Figure 6**).

**Figure 6.**
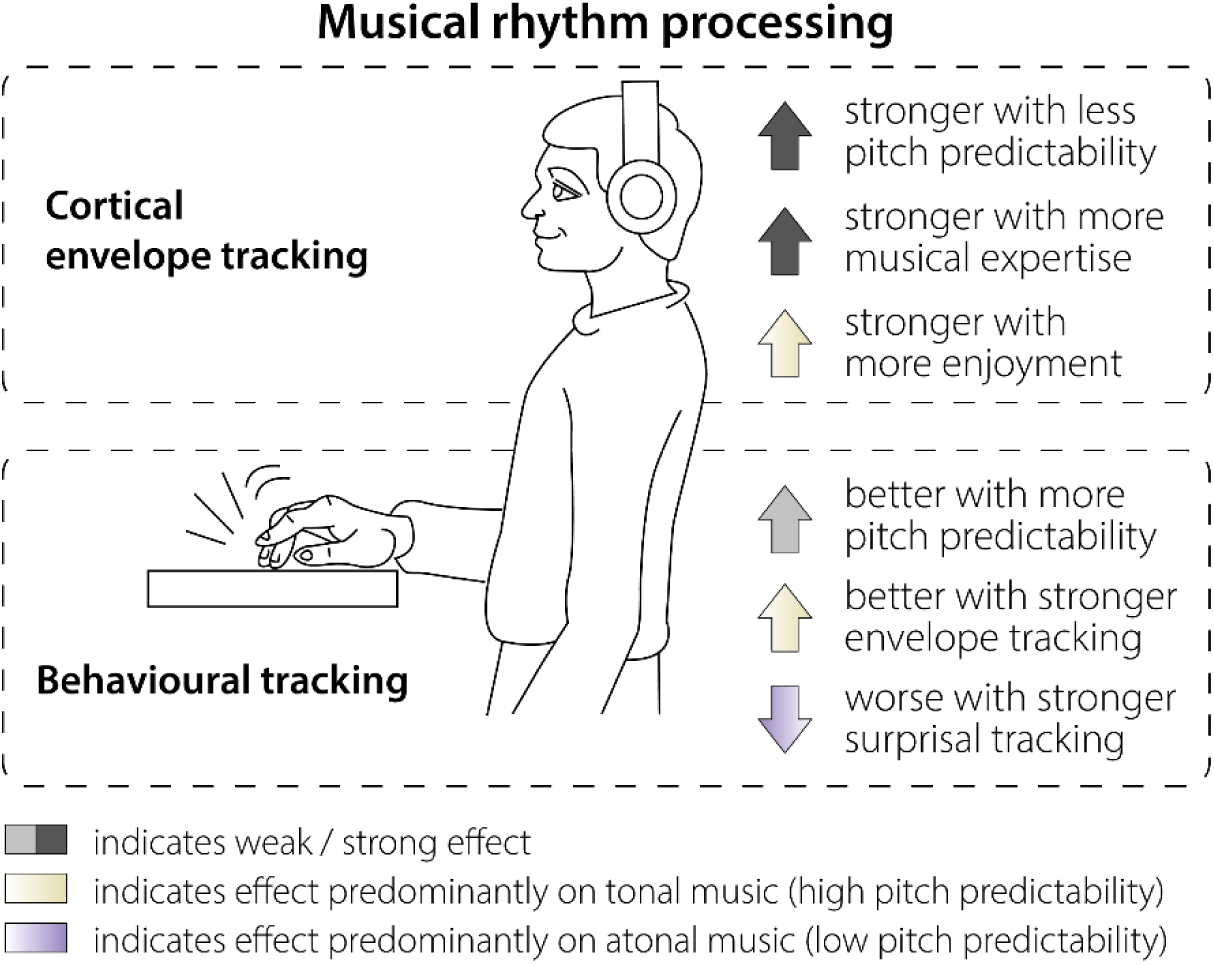
Overview of results.

### Pitch predictability affects rhythm processing as reflected in behavioural tracking

Atonal music can be used to study predictive processing under high uncertainty contexts (Mencke *et al*., 2022; Mencke *et al*., 2018). While temporal predictability has been shown to increase pitch discrimination performance (Herbst & Obleser, 2019; Jones *et al*., 2002), it is unclear whether long-term pitch predictability affects the ability to behaviourally follow the beat particularly in naturalistic musical stimuli. Crucially, in our main experiment and a replication study, we show that when listening to naturalistic music, pitch predictability (modelled on long-term statistics, which reflect exposure to a musical culture, and short-term melodic context) modulates the variability of finger-tapping to the beat. In the tonal condition with higher pitch predictability the finger-tapping performance was more consistent (less variable inter-tap-interval) compared with the atonal low predictability condition. The atonal music in our study contained a rhythmic structure that was identical to the tonal condition, but had generally lower pitch predictability, suggesting that this finding reflects a modulation of behavioural rhythm processing (i.e., finger-tapping to the beat) by pitch predictability. This is in line with and extends previous studies showing expectation effects on musical perception (Herbst & Obleser, 2019; Huron, 2008; Jones *et al*., 2002; Pearce & Wiggins, 2006)

### Cortical tracking of the music envelope in tonal and atonal music

At the neural level, the music envelope was tracked in our study for both tonal and atonal music (Figure 4A). The tracking was observed in both conditions with a centro-temporal topography, in accordance with previous reports suggesting auditory cortex generators of the envelope tracking in speech (Gross et al., 2013; Luo & Poeppel, 2007) and music (Di Liberto et al., 2020b; Doelling & Poeppel, 2015). Some studies reported a right lateralisation for music envelope tracking (Doelling & Poeppel, 2015) in line with the asymmetric sampling in time theory (Poeppel, 2003; Zatorre et al., 2002). The heterogeneous findings in the literature regarding whether a hemispheric lateralisation is observed have been related to various top-down influences (Assaneo et al., 2019; Flinker et al., 2019; Zatorre, 2022).

Atonal music was more strongly tracked at frontal and left parietal electrodes compared with tonal music. Importantly, this was the case, although both conditions had an identical rhythmic structure and there were no significant acoustic differences in the modulation spectrum (**Figure 2D**). We interpret this as evidence that pitch predictability influences neural rhythm tracking. Our control analysis showed that even when the tracking of pitch intervals was partialled out, atonal music was still tracked stronger than tonal music (**Figure S1**). Further, the atonal condition did not carry higher sensory dissonance. These control analyses support the interpretation that it is pitch predictability, and not low-level pitch differences, that affect rhythm tracking. Our results from the neural data are also in line with our behavioural findings in that they suggest an effect of pitch predictability on rhythm processing. Interestingly, Weineck *et al*. (2022) speculated that more predictable music produces stronger neural synchronisation, which our results contradict. However, their paradigm did not manipulate pitch predictability and results are therefore not directly comparable. A predictive coding approach (Friston, 2010; Heilbron & Chait, 2018) could provide a potential explanation for the observed effect: In the atonal condition, notes were generally less predictable than in the tonal condition. In line with the assumption of expectation suppression (Auksztulewicz & Friston, 2016), this likely led to stronger neural prediction errors, which in turn might have resulted in stronger neural responses to the acoustic envelope (not unlike a mismatch-negativity response, e.g., Koelsch *et al*., 2019; Naatanen *et al*., 2007). Accordingly, Kern *et al*. (2022) showed that surprising notes elicit stronger neural responses compared to predictable ones (see also Di Liberto *et al*., 2020a).

### Increased envelope tracking in tonal music related to better tapping performance

In the tonal condition, the cortical tracking of the musical envelope correlated with the behavioural tracking, with higher cortical tracking being associated with an increased ability to behaviourally follow the beat. The findings are in line with previous research showing a positive correlation between behavioural performance and cortical tracking of speech (Schmitt *et al*., 2022; Schubert *et al*., 2023) and music (Doelling & Poeppel, 2015). The relationship between cortical tracking and behavioural performance, however, might be more complex than this, as suggested for speech (Howard & Poeppel, 2010; Pefkou *et al*., 2017). No correlation was observed in the atonal condition and furthermore when the correlation effects were tested in a regression model that included both conditions and selected electrodes, no significant interaction effect was observed. This makes any interpretation regarding differences between tonal and atonal music difficult. A potential explanation is that under conditions of low pitch predictability (as in the atonal condition) the positive relationship between envelope tracking and behavioural tapping performance is confounded, perhaps due to the increased difficulty of trying to (unsuccessfully) make predictions about the upcoming notes. In summary, increased behavioural tracking was related to increased cortical tracking only in the tonal condition, albeit the effect was small.

### Pitch surprisal is cortically tracked in tonal and atonal music

We designed our stimuli so that the pitch predictability of the musical pieces was increased in the atonal compared to the tonal condition (**Figure 5B**). Importantly, our human listeners significantly tracked pitch surprisal in both conditions, which indicates that their pitch expectations (or their prediction errors) were comparable with the IDyOM model. Further, there were no significant condition differences in the cortical tracking of pitch surprisal (**Figure 5A**). This suggests that participants’ neural models of pitch surprisal matches the IDyOM computations, and the notes in the atonal condition elicited not only more surprisal in the IDyOM model, but also in listeners’ neural response. Cortical tracking of pitch surprisal in natural music has been rarely investigated. Two recent studies report melodic surprisal tracking in tonal music that was localised to bilateral superior temporal and Heschl’s gyri (amongst others), and additionally showed either a central topography (using EEG/EcoG, Di Liberto *et al*., 2020a) or a broad fronto-temporal (and central) topography (using MEG, Kern *et al*., 2022). Overall, we found relatively widespread fronto-temporal tracking of pitch surprisal across conditions, which is in line with the above results. We here show that listeners consistently track pitch surprisal not only for high-predictable music, as previously shown, but also for low-predictable music.

### In atonal music lower pitch surprisal tracking is related to better tapping performance

Interestingly, the surprisal tracking strength was only correlated with the behavioural rhythm tracking performance in the atonal but not the tonal condition (as shown by the significant interaction between condition and surprisal tracking). Participants who tracked the pitch surprisal stronger, meaning they matched the high surprisal values from the IDyOM model, also showed more variation (worse performance) in their tapping in the atonal condition. Prediction tendencies have been suggested to vary across participants (Schubert *et al*., 2023). Our measure of surprisal tracking might reflect such a tendency, with some individuals being more or less prone (or able) to make predictions. In the atonal condition with its high uncertainty, pitch predictions might be less informative for rhythm processing, and listeners who tend to make (stronger) predictions, which lead to high prediction errors, could have fewer resources to track the envelope, and to perform well in the tapping task. The few behavioural studies that looked at long-term pitch surprisal tracking have not related it to the rhythm processing performance (e.g., Kern *et al*., 2022). Our results indicate that the negative effect of pitch surprisal tracking on behavioural rhythm processing might only be expected when pitch predictability is low, and prediction errors are high, as in the case of atonal music. The cortical tracking of pitch surprisal was not systematically related to the cortical tracking of the acoustic envelope, at least not in our sample of 20 participants that underwent the EEG part of the study.

### Enjoyment and musical expertise are related to cortical envelope tracking

As expected based on the literature (Mencke *et al*., 2022; Mencke *et al*., 2018), the atonal music condition was rated as less pleasant than the tonal music condition. The individually perceived pleasure or enjoyment has a strong influence on our everyday music listening behaviour (c.f., Gold *et al*., 2019). In the tonal condition, the enjoyment ratings correlated with the strength of cortical music envelope tracking, and this pattern was not statistically different in the atonal condition. The causal nature of this relationship remains unclear: Do listeners show stronger acoustic envelope tracking because they find the music more enjoyable, or do they find it more enjoyable because they have better acoustic envelope tracking? Interestingly, a previous study that investigated whether enjoyment influences neural synchronisation to music did not find a significant effect (Weineck *et al*., 2022). The discrepancy with our findings might be due to differences in experimental paradigms, music choices, quantification of music tracking and analytical methods. If future studies replicate our finding, this suggests that the individual preference of listeners should be taken into account when measuring envelope tracking.

Additionally, musical expertise was correlated positively with the cortical envelope tracking of the music pieces at a cluster of frontal and right temporal electrodes for both the tonal and atonal condition, with more expertise being related to stronger tracking. Our finding is in line with previous reports of effects of musical expertise on cortical envelope tracking (Di Liberto *et al*., 2020b; Doelling & Poeppel, 2015; Harding *et al*., 2019; but see Weineck *et al*., 2022 for a null effect). We here extend these previous results by showing that stronger envelope tracking by individuals with more musical expertise also extends to atonal music with low pitch predictability. The effect of musical expertise on auditory processing of music has been related to increased auditory-motor coupling after musical training (Du & Zatorre, 2017; Rimmele *et al*., 2021).

Another, not mutually exclusive, interpretation for the effects of both enjoyment and expertise, could be that both are associated with generally more attention to the musical stimuli. That is, more musical expertise, as well as more enjoyment of the music, could result in unspecific increases of attention, which in turn could increase the signal-to-noise ratio of acoustic envelope tracking (e.g., Ding & Simon, 2012; Keitel *et al*., 2011; Rimmele *et al*., 2015). To disentangle contributions of attention and other variables, future studies could implement measures of attention or listening effort into their paradigms.

## Conclusion

Our findings suggest that tracking of the envelope of naturalistic music, beyond rhythmic processing, is modulated by top-down factors such as pitch predictability, musical expertise, and enjoyment. In addition to the rhythm, musical pitch surprisal is tracked for both low and high predictable music. This supports the view that long-term musical pitch predictability is processed in the brain and used to facilitate rhythm processing. For tonal, more predictable music, the ability to make valid pitch predictions seems to facilitate the ability to behaviourally follow the rhythm. For atonal music, the reduced pitch predictability results in stronger acoustic envelope tracking than for tonal music, possibly related to higher prediction errors. These higher prediction errors also seem to come at the cost of finger-tapping performance, as individuals with stronger pitch surprisal tracking show worse behavioural rhythm tracking. Overall, our findings indicate that rhythm processing interacts with non-rhythmic stimulus properties, in our case pitch surprisal, and listeners’ characteristics such as music expertise and enjoyment.

## Acknowledgments

AK is supported by the Medical Research Council [grant number MR/W02912X/1]. AK, CK and JR are members of the Scottish-EU Critical Oscillations Network (SCONe), funded by the Royal Society of Edinburgh (RSE Saltire Facilitation Network Award to CK and AK, Reference Number 1963). JR is supported by the Max Planck Institute for Empirical Aesthetics. IM is funded by the Deutsche Forschungsgemeinschaft (DFG, German Research Foundation; project number 510788453). JR, CP and XG are supported by the Max Planck NYU Center for Language, Music, and Emotion (CLaME). We thank Roddy Easson for providing the ‘tapping person’ drawing.

## Supplemental analysis

To explore whether pitch intervals influence acoustic envelope tracking (and our result that envelope tracking is stronger in the atonal than the tonal condition), we performed a conditional MI analysis (Brohl *et al*., 2022; Ince *et al*., 2017). Here, all parameters were identical to the main analysis of acoustic envelope tracking (**Figure 4a**), but we included pitch intervals as the to-be-partialled-out signal in both tonal and atonal conditions.

To extract pitch values, we first used the MIDI toolbox (Eerola & Toiviainen, 2004) for melody and bass lines of the tonal and atonal conditions. The difference between subsequent MIDI pitch values was then used to obtain a measure of pitch intervals, and to create a continuous signal where each pitch interval was as long as the corresponding note (equivalent to how the surprisal signal was created). These pitch interval signals were then normalised using Gaussian Copulas, before including them in the conditional MI analysis. This means each MI computation (for tonal and atonal conditions) included the EEG signal, the envelope signal, and the to-be-partialled out pitch interval signals for both the melody and bass lines.

With pitch intervals in melody and bass lines partialled out, the tracking of the acoustic envelope yielded similar results as the initial tracking analysis (**Figure S1A**). When compared against chance, the envelope was tracked significantly in both tonal and atonal conditions. In the tonal condition, we found a large positive cluster of 31 electrodes that tracked amplitude fluctuations significantly (*p* = .002, *MI*_sum_ = 0.325). Equivalently, in the atonal condition, there was a positive cluster of 32 electrodes that showed significant envelope tracking (*p* < .002, *MI*_sum_ = 0.430). A direct comparison showed that envelope in the atonal condition was tracked stronger than in the tonal condition in two clusters, one over frontal electrodes (*p* = .002, Cohen’s *d*_peak_ = −1.21, 6 electrodes) and one over left occipital electrodes (*p* = .002, Cohen’s *d*_peak_ = −2.17, 3 electrodes). This suggests that, with pitch intervals partialled out, the acoustic envelope is still tracked above chance level, and this tracking is still stronger in the atonal condition.

**Figure S1.**
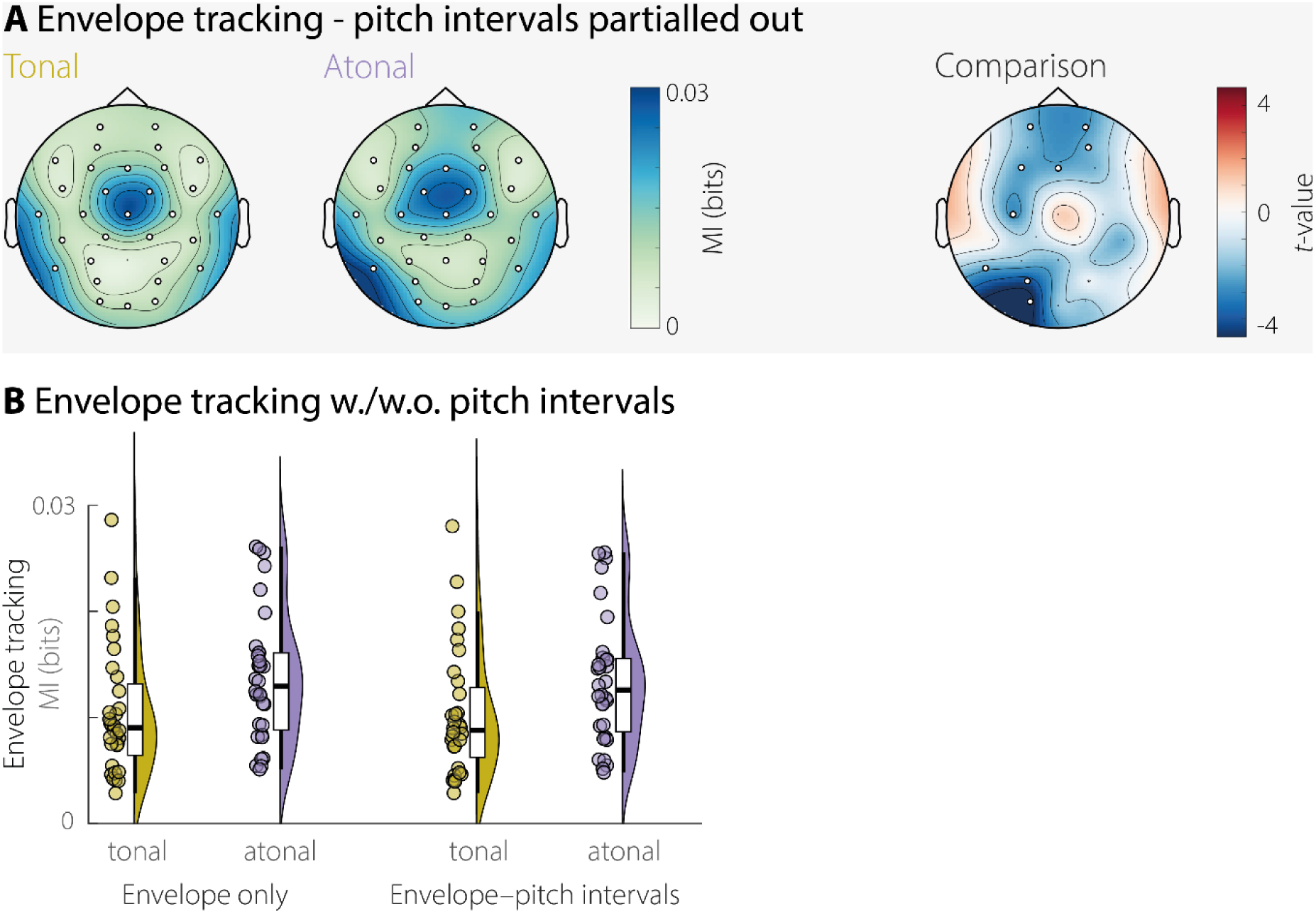
Conditional MI analysis of envelope tracking with pitch intervals partialled out. **A)** Topography of cortical envelope tracking assessed through mutual information (in bits) for both conditions, with partialled out pitch intervals in the melody and bass lines. The right topography shows *t*-values from a direct comparison between tonal and atonal music. **B)** Envelope tracking for all electrodes, averaged across participants, for the tonal and atonal conditions, and for the original MI analysis (tracking of envelope) and the conditional MI analysis (tracking of envelope, pitch intervals partialled out). MI values are systematically lower when pitch intervals are partialled out, but this effect is not different for the tonal or atonal condition.

To directly compare MI values across tracking analyses (with and without pitch intervals partialled out), we averaged MI values for each electrode across participants (**Figure S1B**) and performed a repeated measures 2x2 ANOVA with factors *tracking measure* (envelope tracking vs conditional envelope tracking) and *tonality* (tonal vs atonal). Both main effects were significant (*tracking measure*: *F*(1,31) = 129.93, *p* < .001, η_p_^2^ = .807; *tonality*: *F*(1,31) = 31.39, *p* < .001, η_p_^2^ = .503). The MI values were lower when the tracking of pitch intervals was partialled out (*tonal*: *M*_Env_ = 0.011, SD = 0.006, *M*_Env-Pitch_ = 0.010, SD = 0.006; *atonal*: *M*_Env_ = 0.014, SD = 0.006, *M*_Env-Pitch_ = 0.013, SD = 0.006). MI values were higher in the atonal than the tonal condition, reflecting the difference shown in **Figure S1A**. Crucially, the interaction *tracking measure × tonality* was not significant (*F*(1,31) = 1.51, *p* > .227, η_p_^2^ = .047), which implies that pitch-interval tracking did not affect the conditions differently.

In sum, while there is a small but highly consistent decrease in MI when the tracking of pitch intervals is partialled out, this effect is not different in tonal and atonal conditions. Our previous finding that the acoustic envelope in the atonal condition is tracked stronger than in the tonal condition, therefore also holds true in the conditional MI analysis. This means that the found differences in envelope tracking are unlikely to be due to low-level differences in the tracking of pitch intervals.

## References

Abel, S., Dressel, K., Bitzer, R., Kummerer, D., Mader, I., Weiller, C., & Huber, W. (2009). The separation of processing stages in a lexical interference fMRI-paradigm. Neuroimage, 44(3), 1113–1124. 10.1016/j.neuroimage.2008.10.018

Abrams, E. B., Vidal, E. M., Pelofi, C., & Ripollés, P. (2022). Retrieving musical information from neural data: how cognitive features enrich acoustic ones. Ismir 2022 Hybrid Conference,

Alexandrou, A., Saarinen, T., Kujala, J., & Salmelin, R. (2018). Cortical entrainment: what we can learn from studying naturalistic speech perception. Language, Cognition and Neuroscience, 1–13.

Arachchige, C., Prendergast, L. A., & Staudte, R. G. (2022). Robust analogs to the coefficient of variation. J Appl Stat, 49(2), 268–290. 10.1080/02664763.2020.1808599

Assaneo, M. F., Rimmele, J. M., Orpella, J., Ripolles, P., de Diego-Balaguer, R., & Poeppel, D. (2019). The Lateralization of Speech-Brain Coupling Is Differentially Modulated by Intrinsic Auditory and Top-Down Mechanisms. Front Integr Neurosci, 13, 28. 10.3389/fnint.2019.00028

Auksztulewicz, R., & Friston, K. (2016). Repetition suppression and its contextual determinants in predictive coding. cortex, 80, 125–140. 10.1016/j.cortex.2015.11.024

Benjamini, Y., & Hochberg, Y. (1995). Controlling the False Discovery Rate - a Practical and Powerful Approach to Multiple Testing. Journal of the Royal Statistical Society Series B-Methodological, 57(1), 289–300. <Go to ISI>://WOS:A1995QE45300017

Blanco-Elorrieta, E., Ding, N., Pylkkänen, L., & Poeppel, D. (2020). Understanding requires tracking: noise and knowledge interact in bilingual comprehension. Journal of cognitive neuroscience, 32(10), 1975–1983.

Brohl, F., & Kayser, C. (2021). Delta/theta band EEG differentially tracks low and high frequency speech-derived envelopes. Neuroimage, 233, 117958. 10.1016/j.neuroimage.2021.117958

Brohl, F., Keitel, A., & Kayser, C. (2022). MEG Activity in Visual and Auditory Cortices Represents Acoustic Speech-Related Information during Silent Lip Reading. eNeuro, 9(3). 10.1523/ENEURO.0209-22.2022

Celma-Miralles, A., & Toro, J. M. (2019). Ternary meter from spatial sounds: Differences in neural entrainment between musicians and non-musicians. Brain Cogn, 136, 103594. 10.1016/j.bandc.2019.103594

Cervantes Constantino, F., & Simon, J. Z. (2018). Restoration and efficiency of the neural processing of continuous speech are promoted by prior knowledge. Frontiers in systems neuroscience, 12, 56.

Cuddy, L. L., Cohen, A. J., & Mewhort, D. J. (1981). Perception of structure in short melodic sequences. J Exp Psychol Hum Percept Perform, 7(4), 869–883. 10.1037//0096-1523.7.4.869

Di Liberto, G. M., Lalor, E. C., & Millman, R. E. (2018). Causal cortical dynamics of a predictive enhancement of speech intelligibility. Neuroimage, 166, 247–258.

Di Liberto, G. M., Pelofi, C., Bianco, R., Patel, P., Mehta, A. D., Herrero, J. L., Mesgarani, N. (2020a). Cortical encoding of melodic expectations in human temporal cortex. Elife, 9, e51784.

Di Liberto, G. M., Pelofi, C., Shamma, S., & de Cheveigné, A. (2020b). Musical expertise enhances the cortical tracking of the acoustic envelope during naturalistic music listening. Acoustical Science and Technology, 41(1), 361–364.

Dibben, N. (1994). The cognitive reality of hierarchic structure in tonal and atonal music. Music Perception, 12(1), 1–25.

Ding, N., Patel, A. D., Chen, L., Butler, H., Luo, C., & Poeppel, D. (2017). Temporal modulations in speech and music. Neurosci Biobehav Rev, 81(Pt B), 181–187. 10.1016/j.neubiorev.2017.02.011

Ding, N., & Simon, J. Z. (2012). Emergence of neural encoding of auditory objects while listening to competing speakers. Proc Natl Acad Sci U S A, 109(29), 11854–11859. 10.1073/pnas.1205381109

Doelling, K. B., & Poeppel, D. (2015). Cortical entrainment to music and its modulation by expertise. Proceedings of the National Academy of Sciences, 112(45), E6233–E6242.

Dowling, W. J., Kwak, S., & Andrews, M. W. (1995). The time course of recognition of novel melodies. Perception & Psychophysics, 57, 136–149.

Du, Y., & Zatorre, R. J. (2017). Musical training sharpens and bonds ears and tongue to hear speech better. Proc Natl Acad Sci U S A, 114(51), 13579–13584. 10.1073/pnas.1712223114

Engel, A. K., Fries, P., & Singer, W. (2001). Dynamic predictions: oscillations and synchrony in top-down processing. Nat Rev Neurosci, 2(10), 704–716. 10.1038/35094565

Fiveash, A., Bella, S. D., Bigand, E., Gordon, R. L., & Tillmann, B. (2022). You got rhythm, or more: The multidimensionality of rhythmic abilities. Atten Percept Psychophys, 84(4), 1370–1392. 10.3758/s13414-022-02487-2

Flinker, A., Doyle, W. K., Mehta, A. D., Devinsky, O., & Poeppel, D. (2019). Spectrotemporal modulation provides a unifying framework for auditory cortical asymmetries. Nat Hum Behav, 3(4), 393–405. 10.1038/s41562-019-0548-z

Friston, K. (2010). The free-energy principle: a unified brain theory? Nat Rev Neurosci, 11(2), 127–138. 10.1038/nrn2787

Gold, B. P., Pearce, M. T., Mas-Herrero, E., Dagher, A., & Zatorre, R. J. (2019). Predictability and uncertainty in the pleasure of music: a reward for learning? Journal of Neuroscience, 39(47), 9397–9409.

Gorsuch, R. L., & Lehmann, C. S. (2010). Correlation coefficients: Mean bias and confidence interval distortions. Journal of Methods and Measurement in the Social Sciences, 1(2), 52–65.

Gross, J., Hoogenboom, N., Thut, G., Schyns, P., Panzeri, S., Belin, P., & Garrod, S. (2013). Speech rhythms and multiplexed oscillatory sensory coding in the human brain. PLoS Biol, 11(12), e1001752. 10.1371/journal.pbio.1001752

Guan, X., Ren, Z., & Pelofi, C. (2022). py2lispIDyOM: A Python package for the information dynamics of music (IDyOM) model. Journal of Open Source Software, 7(79), 4738.

Harding, E. E., Sammler, D., Henry, M. J., Large, E. W., & Kotz, S. A. (2019). Cortical tracking of rhythm in music and speech. Neuroimage, 185, 96–101.

Heilbron, M., & Chait, M. (2018). Great Expectations: Is there Evidence for Predictive Coding in Auditory Cortex? Neuroscience, 389, 54–73. 10.1016/j.neuroscience.2017.07.061

Herbst, S. K., & Obleser, J. (2019). Implicit temporal predictability enhances pitch discrimination sensitivity and biases the phase of delta oscillations in auditory cortex. Neuroimage, 203, 116198. 10.1016/j.neuroimage.2019.116198

Howard, M. F., & Poeppel, D. (2010). Discrimination of speech stimuli based on neuronal response phase patterns depends on acoustics but not comprehension. J Neurophysiol, 104(5), 2500–2511. 10.1152/jn.00251.2010

Huron, D. (2008). Sweet anticipation: Music and the psychology of expectation. MIT press.

Ince, R. A., Giordano, B. L., Kayser, C., Rousselet, G. A., Gross, J., & Schyns, P. G. (2017). A statistical framework for neuroimaging data analysis based on mutual information estimated via a gaussian copula. Hum Brain Mapp, 38(3), 1541–1573. 10.1002/hbm.23471

Iversen, J. R., Patel, A. D., Nicodemus, B., & Emmorey, K. (2015). Synchronization to auditory and visual rhythms in hearing and deaf individuals. Cognition, 134, 232–244. 10.1016/j.cognition.2014.10.018

Jones, M. R., Moynihan, H., MacKenzie, N., & Puente, J. (2002). Temporal aspects of stimulus-driven attending in dynamic arrays. Psychol Sci, 13(4), 313–319. 10.1111/1467-9280.00458

Keitel, A., Gross, J., & Kayser, C. (2018). Perceptually relevant speech tracking in auditory and motor cortex reflects distinct linguistic features. PLoS Biol, 16(3), e2004473.

Keitel, A., Gross, J., & Kayser, C. (2020). Shared and modality-specific brain regions that mediate auditory and visual word comprehension. Elife, 9. 10.7554/eLife.56972

Keitel, A., Ince, R. A., Gross, J., & Kayser, C. (2017). Auditory cortical delta-entrainment interacts with oscillatory power in multiple fronto-parietal networks. Neuroimage, 147, 32–42. 10.1016/j.neuroimage.2016.11.062

Keitel, C., Obleser, J., Jessen, S., & Henry, M. J. (2021). Frequency-Specific Effects in Infant Electroencephalograms Do Not Require Entrained Neural Oscillations: A Commentary on Koster et al. (2019). Psychol Sci, 32(6), 966–971. 10.1177/09567976211001317

Keitel, C., Schroger, E., Saupe, K., & Muller, M. M. (2011). Sustained selective intermodal attention modulates processing of language-like stimuli. Exp Brain Res, 213(2-3), 321–327. 10.1007/s00221-011-2667-2

Kern, P., Heilbron, M., de Lange, F. P., & Spaak, E. (2022). Cortical activity during naturalistic music listening reflects short-range predictions based on long-term experience. eLife, 11. 10.7554/eLife.80935

Koelsch, S., Vuust, P., & Friston, K. (2019). Predictive Processes and the Peculiar Case of Music. Trends Cogn Sci, 23(1), 63–77. 10.1016/j.tics.2018.10.006

Koike, K. J., Hurst, M. K., & Wetmore, S. J. (1994). Correlation between the American-Academy-of-Otolaryngology-Head-and-Neck-Surgery 5-minute hearing test and standard audiological data. Otolaryngology-Head and Neck Surgery, 111(5), 625–632. <Go to ISI>://A1994PR79400014

Lakens, D. (2013). Calculating and reporting effect sizes to facilitate cumulative science: a practical primer for t-tests and ANOVAs. Front Psychol, 4, 863. 10.3389/fpsyg.2013.00863

Lartillot, O., Toiviainen, P., & Eerola, T. (2008). A Matlab Toolbox for Music Information Retrieval. In C. Preisach, H. Burkhardt, L. Schmidt-Thieme, & R. Decker (Eds.), Data Analysis, Machine Learning and Applications. Springer-Verlag.

Lerdahl, F. (2019). Composition and cognition: Reflections on contemporary music and the musical mind. University of California Press.

Lesenfants, D., & Francart, T. (2020). The interplay of top-down focal attention and the cortical tracking of speech. Scientific reports, 10(1), 1–10.

Leys, C., Ley, C., Klein, O., Bernard, P., & Licata, L. (2013). Detecting outliers: Do not use standard deviation around the mean, use absolute deviation around the median. Journal of experimental social psychology, 49(4), 764–766.

Luo, H., & Poeppel, D. (2007). Phase patterns of neuronal responses reliably discriminate speech in human auditory cortex. Neuron, 54(6), 1001–1010. 10.1016/j.neuron.2007.06.004

Marion, G., Di Liberto, G. M., & Shamma, S. A. (2021). The Music of Silence: Part I: Responses to Musical Imagery Encode Melodic Expectations and Acoustics. Journal of Neuroscience, 41(35), 7435–7448.

Mencke, I., Omigie, D., Quiroga-Martinez, D. R., & Brattico, E. (2022). Atonal Music as a Model for Investigating Exploratory Behavior. Frontiers in Neuroscience, 16, 793163.

Mencke, I., Omigie, D., Wald-Fuhrmann, M., & Brattico, E. (2018). Atonal Music: Can Uncertainty Lead to Pleasure? Front Neurosci, 12, 979. 10.3389/fnins.2018.00979

Mencke, I., Quiroga-Martinez, D. R., Omigie, D., Michalareas, G., Schwarzacher, F., Haumann, N. T., Brattico, E. (2021). Prediction under uncertainty: Dissociating sensory from cognitive expectations in highly uncertain musical contexts. Brain Res, 1773, 147664. 10.1016/j.brainres.2021.147664

Morillon, B., Arnal, L. H., Schroeder, C. E., & Keitel, A. (2019). Prominence of delta oscillatory rhythms in the motor cortex and their relevance for auditory and speech perception. Neuroscience & Biobehavioral Reviews. 10.1016/j.neubiorev.2019.09.012

Naatanen, R., Paavilainen, P., Rinne, T., & Alho, K. (2007). The mismatch negativity (MMN) in basic research of central auditory processing: a review. Clin Neurophysiol, 118(12), 2544–2590. 10.1016/j.clinph.2007.04.026

Neuloh, G., & Curio, G. (2004). Does familiarity facilitate the cortical processing of music sounds? Neuroreport, 15(16), 2471–2475. 10.1097/00001756-200411150-00008

Nozaradan, S., Peretz, I., & Mouraux, A. (2012). Selective neuronal entrainment to the beat and meter embedded in a musical rhythm. J Neurosci, 32(49), 17572–17581. 10.1523/JNEUROSCI.3203-12.2012

Obleser, J., & Kayser, C. (2019). Neural Entrainment and Attentional Selection in the Listening Brain. Trends in cognitive sciences, 23(11), 913–926. 10.1016/j.tics.2019.08.004

Ockelford, A., & Sergeant, D. (2013). Musical expectancy in atonal contexts: Musicians’ perception of “antistructure”. Psychology of Music, 41(2), 139–174.

Oldfield, R. C. (1971). The assessment and analysis of handedness: the Edinburgh inventory. Neuropsychologia, 9(1), 97–113. http://www.ncbi.nlm.nih.gov/pubmed/5146491

Oostenveld, R., Fries, P., Maris, E., & Schoffelen, J. M. (2011). FieldTrip: Open source software for advanced analysis of MEG, EEG, and invasive electrophysiological data. Comput Intell Neurosci, 2011, 156869. 10.1155/2011/156869

Park, H., & Kayser, C. (2019). Shared neural underpinnings of multisensory integration and trial-by-trial perceptual recalibration in humans. Elife, 8.

Pearce, M. T. (2005). The construction and evaluation of statistical models of melodic structure in music perception and composition https://openaccess.city.ac.uk/id/eprint/8459

Pearce, M. T. (2018). Statistical learning and probabilistic prediction in music cognition: mechanisms of stylistic enculturation. Ann N Y Acad Sci, 1423(1), 378–395. 10.1111/nyas.13654

Pearce, M. T., Ruiz, M. H., Kapasi, S., Wiggins, G. A., & Bhattacharya, J. (2010). Unsupervised statistical learning underpins computational, behavioural, and neural manifestations of musical expectation. Neuroimage, 50(1), 302–313.

Pearce, M. T., & Wiggins, G. A. (2006). Expectation in Melody: The Influence of Context and Learning. Music Perception, 23(5), 377–405. 10.1525/mp.2006.23.5.377

Pearce, M. T., & Wiggins, G. A. (2012). Auditory expectation: the information dynamics of music perception and cognition. Top Cogn Sci, 4(4), 625–652. 10.1111/j.1756-8765.2012.01214.x

Peelle, J. E., Gross, J., & Davis, M. H. (2013). Phase-locked responses to speech in human auditory cortex are enhanced during comprehension. Cereb Cortex, 23(6), 1378–1387. 10.1093/cercor/bhs118

Pefkou, M., Arnal, L. H., Fontolan, L., & Giraud, A. L. (2017). theta-band and beta-band neural activity reflects independent syllable tracking and comprehension of time-compressed speech. J Neurosci, 37(33), 7930–7938. 10.1523/JNEUROSCI.2882-16.2017

Poeppel, D. (2003). The analysis of speech in different temporal integration windows: cerebral lateralization as ‘asymmetric sampling in time’. Speech Communication, 41(1), 245–255. 10.1016/S0167-6393(02)00107-3

Reetzke, R., Gnanateja, G. N., & Chandrasekaran, B. (2021). Neural tracking of the speech envelope is differentially modulated by attention and language experience. Brain Lang, 213, 104891. 10.1016/j.bandl.2020.104891

Repp, B. H. (2005). Sensorimotor synchronization: a review of the tapping literature. Psychon Bull Rev, 12(6), 969–992. 10.3758/bf03206433

Rimmele, J. M., Kern, P., Lubinus, C., Frieler, K., Poeppel, D., & Assaneo, M. F. (2021). Musical Sophistication and Speech Auditory-Motor Coupling: Easy Tests for Quick Answers. Front Neurosci, 15, 764342. 10.3389/fnins.2021.764342

Rimmele, J. M., Zion Golumbic, E., Schroger, E., & Poeppel, D. (2015). The effects of selective attention and speech acoustics on neural speech-tracking in a multi-talker scene. Cortex, 68, 144–154. 10.1016/j.cortex.2014.12.014

Rovetti, J., Copelli, F., & Russo, F. A. (2022). Audio and visual speech emotion activate the left pre-supplementary motor area. Cogn Affect Behav Neurosci, 22(2), 291–303. 10.3758/s13415-021-00961-2

Sammler, D. (2020). Splitting speech and music. science, 367(6481), 974–976. 10.1126/science.aba7913

Schmitt, R., Meyer, M., & Giroud, N. (2022). Better speech-in-noise comprehension is associated with enhanced neural speech tracking in older adults with hearing impairment. cortex, 151, 133–146. 10.1016/j.cortex.2022.02.017

Schubert, J., Schmidt, F., Gehmacher, Q., Bresgen, A., & Weisz, N. (2023). Cortical speech tracking is related to individual prediction tendencies. Cereb Cortex, 33(11), 6608–6619. 10.1093/cercor/bhac528

Schulze, K., Jay Dowling, W., & Tillmann, B. (2011). Working memory for tonal and atonal sequences during a forward and a backward recognition task. Music Perception, 29(3), 255–267.

Smith, Z. M., Delgutte, B., & Oxenham, A. J. (2002). Chimaeric sounds reveal dichotomies in auditory perception. Nature, 416(6876), 87–90. 10.1038/416087a

Song, J., & Iverson, P. (2018). Listening effort during speech perception enhances auditory and lexical processing for non-native listeners and accents. Cognition, 179, 163–170.

Tierney, A., & Kraus, N. (2015). Neural entrainment to the rhythmic structure of music. J Cogn Neurosci, 27(2), 400–408. 10.1162/jocn_a_00704

Vuvan, D. T., Podolak, O. M., & Schmuckler, M. A. (2014). Memory for musical tones: the impact of tonality and the creation of false memories. Front Psychol, 5, 582. 10.3389/fpsyg.2014.00582

Weineck, K., Wen, O. X., & Henry, M. J. (2022). Neural synchronization is strongest to the spectral flux of slow music and depends on familiarity and beat salience. eLife, 11. 10.7554/eLife.75515

Zarate, J. M., Tian, X., Woods, K. J., & Poeppel, D. (2015). Multiple levels of linguistic and paralinguistic features contribute to voice recognition. Sci Rep, 5, 11475. 10.1038/srep11475

Zatorre, R. J. (2001). Neural specializations for tonal processing. Ann N Y Acad Sci, 930, 193–210. 10.1111/j.1749-6632.2001.tb05734.x

Zatorre, R. J. (2022). Hemispheric asymmetries for music and speech: Spectrotemporal modulations and top-down influences. Front Neurosci, 16, 1075511. 10.3389/fnins.2022.1075511

Zatorre, R. J., Belin, P., & Penhune, V. B. (2002). Structure and function of auditory cortex: music and speech. Trends Cogn Sci, 6(1), 37–46. http://www.ncbi.nlm.nih.gov/pubmed/11849614

Zion Golumbic, E. M., Ding, N., Bickel, S., Lakatos, P., Schevon, C. A., McKhann, G. M., . . . Schroeder, C. E. (2013). Mechanisms underlying selective neuronal tracking of attended speech at a “cocktail party”. Neuron, 77(5), 980–991. 10.1016/j.neuron.2012.12.037

Zuk, N. J., Murphy, J. W., Reilly, R. B., & Lalor, E. C. (2021). Envelope reconstruction of speech and music highlights stronger tracking of speech at low frequencies. PLoS Comput Biol, 17(9), e1009358. 10.1371/journal.pcbi.1009358

## References

Eerola, T., & Toiviainen, P. (2004). MIDI Toolbox: MATLAB Tools for Music Research. University of Jyväskylä.

